# An application of higher order multivariate cumulants in modelling of myoelectrical activity of porcine uterus during early pregnancy

**DOI:** 10.1101/443564

**Authors:** Malgorzata Domino, Krzysztof Domino, Zdzislaw Gajewski

## Abstract

The analysis of the uterine contraction have become a general practice in an effort to improve the clinical management of uterine contractions during pregnancy and labour in human beings. The fluctuations in uterine activity may occur without affecting progress of gestation, however the painful and fashion contractions may be the first threat of miscarriage. While pigs were considered as an referential preclinical model, the computational modelling of spontaneous myoelectrical activity of complex systems of porcine myometrium in peri-fertilization period has been proposed. The higher order statistic, multivariate cumulants and Joint Skewness Band Selection method, have been applied to study the dependence structure of electromyographic (EMG) signal with an effective EMG feature. Than the model of recognition of multivariate, myoelectricaly changes according to crucial stages for successful fertilization and early pregnancy maintenance has been estimated. We found that considering together time and frequency features of EMG signal was extremely non-Gaussian distributed and the higher order multivariate statistics such as cumulants, have to be used to determine the pattern of myoelectrical activity in reproductive tract. We confirmed the expectance that the probabilistic model changes on a daily base. We demonstrated the changes in proposed model at the crucial time points of in peri-fertilization period. We speculate the activity of the middle of uterine horn and the power (minimum and maximum) and pauses between myoelectrical burst features are essential for the functional role of uterine contractility in peri-fertilization period.

## 1. Introduction

The analysis of the uterine contraction have become a general practice in an effort to improve the clinical management of uterine contractions during pregnancy and labour (Figueroa et al., 1987; Gajewski and Faundez, 1992; Maul et al., 2003; Rabotti et al., 2008; Jacod et al., 2010; Lammers, 2013; Rooijakkers et al., 2014; Pawlinski et al., 2017). In healthy, no pregnant uteri specific contractile patterns evolve during the estrus cycle both in humans and animals (Fanchin and Ayoubi, 2009; Kuijsters et al., 2017; Domino et. al, 2018a). Significant attention has been dedicated to evolving patterns being in line with the hypothesis of a functional role of uterine activity in promotion of fertilization (Pusey et al., 1980; Sammali et al., 2018). Uterine contractility is known to affect embryo implantation (Rogers et al., 1983; Ziecik et al., 2011; Lammers, 2013). Furthermore, the quiescence of the uterus during pregnancy is required to maintain pregnancy and allow adequate nourishment and development of the foetus (Gajewski and Faundez, 1992; Rabboti and Mischi, 2015; Markiewicz et al., 2016). During early pregnancy, some fluctuations in uterine activity can always be present without affecting progress of gestation, however the painful uterine contractions occurring in a coordinated and forceful fashion may be the first threat of a preterm delivery (Norwitz and Robinson, 2001).

The animal and *in vitro* experiments have been found to provide valuable methods and results for parameter estimation, undertaking up to humans clinical studies (Devedeux et al., 1993; Rabotti et al., 2010). While pigs are considered as an ideal preclinical model due to many anatomical and functional similarities with humans, the investigations carried out on the porcine reproductive tract may be treated as a referential (Ziecik et al., 1993; Gajewski et al, 2001; Kobayashi et al., 2012). The electrical activity of the porcine uterus as the primary cause of contractions, was established both during estrus cycle (Gajewski et al., 2001; Domino et al., 2018a) and early pregnancy (Maner et al., 2003; Markiewicz et al., 2016; Pawlinski et al., 2017). In those cases smooth muscle cells (SMCs) contract, action potentials reach a depolarization threshold and generate an electromagnetic field (Eswaran et al., 2004). The billions of SMCs are coupled electrically by the gap junctions into the complex biological system (Garfield et al., 1977) within numerous electrochemical events generate electric currents as a sum of action potential differences between SMCs (Alkan and Günay, 2012). Those myometrial electromagnetic field is possible to be measured as voltage and quantified with different sensitivity by electromyography (EMG) (Gajewski et al., 2001; Wolinski et al., 2003; Eswaran et al., 2004) and electrohysterography (EHG) (Rabotti et al., 2008; Jacod et al., 2010; Rabotti et al., 2010; Sammali et al., 2018). The EHG was reported as a feasible option to evaluate noninvasively and objectively the myometrial activity in humans, both of the non-pregnant uterus (Sammali et al., 2018) as well as during late gestation and labour (Jacod et al., 2010; Rabotti et al., 2010). In early gestation in humans, EHG method is insufficient to register the subtle changes in spectral content of uterine EMG signal, whereas invasive EMG method is unacceptable for obvious, ethical reasons (Rabotti et al., 2010). The needle EMG gathered the bioelectrical signal directly associated with the contractile activity of myometrium, while EHG collected signal abundant in noise generated between uterus and implanted superficial electrodes positioned on the surface of abdomen (Devedeux et al., 1993; Jacod et al., 2010). Therefore, the electrical activity recorded more invasively, directly from myometrium, may provide crucial information about uterine activity. For this reason the animal models are essential for understanding the sophisticated mechanisms behind the phenomena of contribution of uterine activity into the fertilization and early pregnancy maintenance.

In the case of all electroencephalograms (EEG) (Singh et al., 2015; Chen et al., 2016), EMG (Domino et al., 2016; 2017a; 2018a) and EHG (Farina and Negro, 2007; Garfield and Maner, 2007a; Domino et al., 2017a), considering spectral content in addition to studying changes in its time domain features is widely used. Moreover a variety of features have been extracted by different feature selection methods in the time and frequency domains. The time domain features of EMG include duration (D), amplitude (A) and root mean square (RMS) of bursts as well as duration of pauses between bursts (P) (Devedeux et al., 1993; Gajewski et al., 2004; Garfield et al., 2007a; Pawlinski et al., 2016). It has been established that the uterine EMG signals can be quantified sufficiently with mathematical functions and transforms such as power spectral analysis (Maner et al., 2003; Garfield and Maner, 2007b). In frequency domain we receive features corresponding to each frequency bands. However in our work for simplicity and informative significance we concentrate on: dominant frequency (DF), maximum power (Max P) and the minimum power (Min P) which can be estimated by Fast Fourier transform (FFT) method (Oczeretko et al., 2007; Domino et al., 2016; 2017a; 2018a). Recent methods used in a detailed analysis of myoelectrical signals are divided in four major categories: cross-coherence function (Farina and Merletti, 2004; Domino et al., 2016), phase difference (Hunter et al., 1987), maximum likelihood (Farina et al., 2001; Rabotti et al., 2010) and the detection of spectral relations (Farina and Negro, 2007; Domino et al., 2017; Garfield and Maner 2007b). Garfield and Maner (2007b) suggested the features of power density spectrum as a more effective than the conventional time dependent features in reflecting the actual function of the uterus. The decomposition of myoelectrical signals into their individual frequency or ‘scale’ subcomponents provided much more advantageous data, that can be used as indicators of changes in uterine status such as pre-term delivery (Maner et al., 2003; Garfield and Maner, 2007b). In our research we have acquired both recent features (D, A, RMS, P) and following frequency domain subcomponents (DF, Max P, Min P) of EMG signal, from porcine uterus in relation to early pregnancy phenomena. Further we speculate that those features are non-Gaussian distributed due to complex biochemical processes that take place in myometrium, which represents contractile element of the uterine wall composing of SMCs (Rabboti and Mischi, 2015). Observe that higher order multivariate cumulants can be used to analyse non-Gaussian distributed multivariate data (Domino et al., 2018b) such as considering together time and frequency features in our case. Observe as well, that higher order cumulants approach was successfully applied in EEG (Becker et al., 2014) and superficial EMG (Ju and Liu, 2011) data analysis. However, none of recent researchers has introduced the computational modelling of spontaneous myoelectrical activity of complex SMCs systems in peri-fertilization period.

## 2. Materials and methods

### 2.1. Experimental design

The spontaneous myoelectrical activity of uterus has been recorded from 8 Polish Landrace sows according to protocol approved by the III Local Ethical Committee on Animal Testing in Warsaw (Permit Number: 71/2009, from 19.11.2009) on behalf of the National Ethical Committees on Animal Testing. The study was conducted on sexually mature sows at the age of 6 months and 95–110 kg body weight. All sows were selected from pig producing units after experiencing a single estrous cycle and had not been inseminated. Sows were adapted to the animal facilities for 7 days before beginning the studies and during the entire experiment were housed in experimental cages, fed and watered *ad libitum*. The implantable telemetry method has been applied to obtain the EMG signal from subsequent parts of corpus uteri and unilateral uterine horn. The experiment has been started on non-pregnant sows during diestrus, subsequently the induction and synchronization of estrus was performed to manage an artificial insemination (AI) according to Pawlinski et al. (2017) (day −4: eCG, 750 IU/animal, i.m., Werfaser, Fa; Alvetra, Austria; day −1: hCG, 500 IU/animal, i.m., Werfaser, Fa; Alvetra, Austria; day 0: AI). Indeed, spontaneous uterine activity was registered during long-term registration both in non-pregnant and pregnant state since day 12 after AI.

The surgery under general anesthesia has been performed in order to position the combination of three silver, bipolar, needle electrodes connected to 3-channel telemetry transmitter TL10M3-D70-EEE (DSI, USA) according to Gajewski et al. (2004). Animals were premedicated with an azaperone (Stresnil, 3.0 mg/kg b.w., i.m., Fa; Janssen Pharmaceutica, Belgium). General anesthesia was conducted using combined administration of medetomidine (Cepetor, 1.0 mg/kg b.w., i.v., Fa; CP-Pharma HgmbH, Germany), butorphanol (Butomidor, 0.2 mg/kg b.w., i.v., Fa; Richter Pharma AG, Austria), ketamine (Bioketan, 3.0 mg/kg b.w., i.v., Fa: Vetoquinol Biowet, Poland) and propofol (Propofol, 2.0-4.0 mg/kg b.wt., i.v., Fa; Pfizer, USA). After surgery meloxicam (Metacam, 0.4 mg/kg b.w., i.m., Fa; Boehringer Ingelheim, Germany) and cefquinom (Cobactan, 2.0 mg/kg b.w., i.m., Fa. Intervet, Poland) had been administered for five days. The electrodes were sutured unilaterally onto the surfaces of uterus: the tip of right uterine horn, (channel 1), the middle of right uterine horn (channel 2) and corpus uteri (channel 3). Channel 1 was positioned at a distance of 2 cm from oviduct, whereas distance between the other two channels was fixed at 30 cm. The 3-channel telemetry transmitter was positioned between abdominal muscles directly under the skin (Fig. 1A). Finally, the silicone cannula was inserted into the brachial vein in order to daily blood sample collection.

**Fig. 1.**
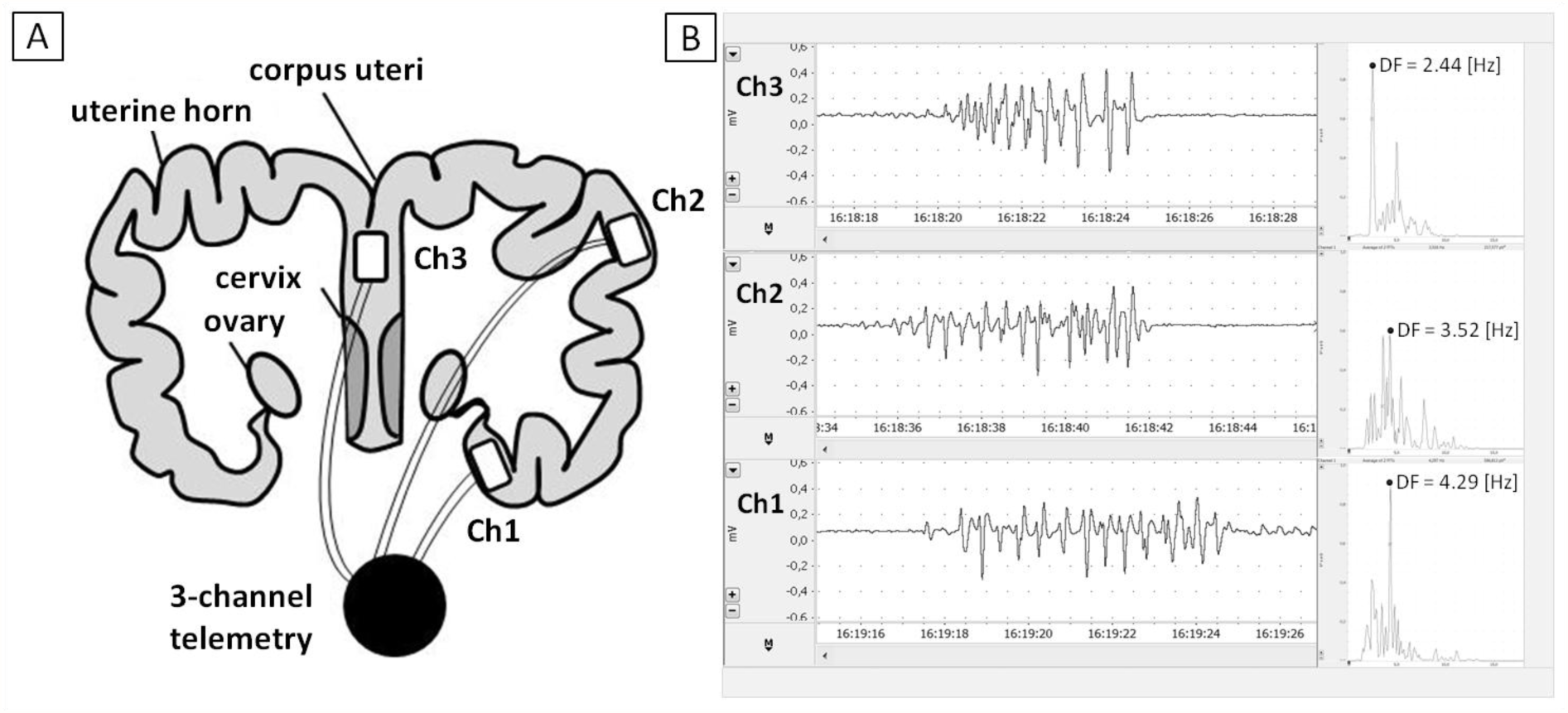
A. The unilateral arranged of the electrodes (Ch1, Ch2, Ch3) in the different parts of uterus. B. The samples of EMG signal in time and frequency domain obtained at day 0 in Ch1, Ch2, Ch3, respectively.

#### 2.1.1 Time-line determination

The experimental time-line was determined based on the hormonal and ovarian status as well as the time of AI (day 0) (Fig. 2). The daily blood samples were used for hormones concentrations: luteinizing hormones (LH) and progesterone (P_4_), determination. Blood samples (7.0 ml every 4 h) collected into vacutainer tubes (Fa; KABE, Poland) were centrifuged (5.590 g; 10 min) and the serum samples were frozen until assaying. The hormones concentrations was determined using radioimmunoassay: a high sensitivity, noncommercial for LH (intra-assay coefficient of variation <6.7% and inter-assay coefficient of variation <12.5%, Ziecik et al., 1978) and a commercial for P_4_ (intra-assay coefficient of variation <5.6% and inter-assay coefficient of variation <8.8%; KIP 1458; Fa; DIAsource ImmunoAssays SA, Belgium).

**Fig. 2.**
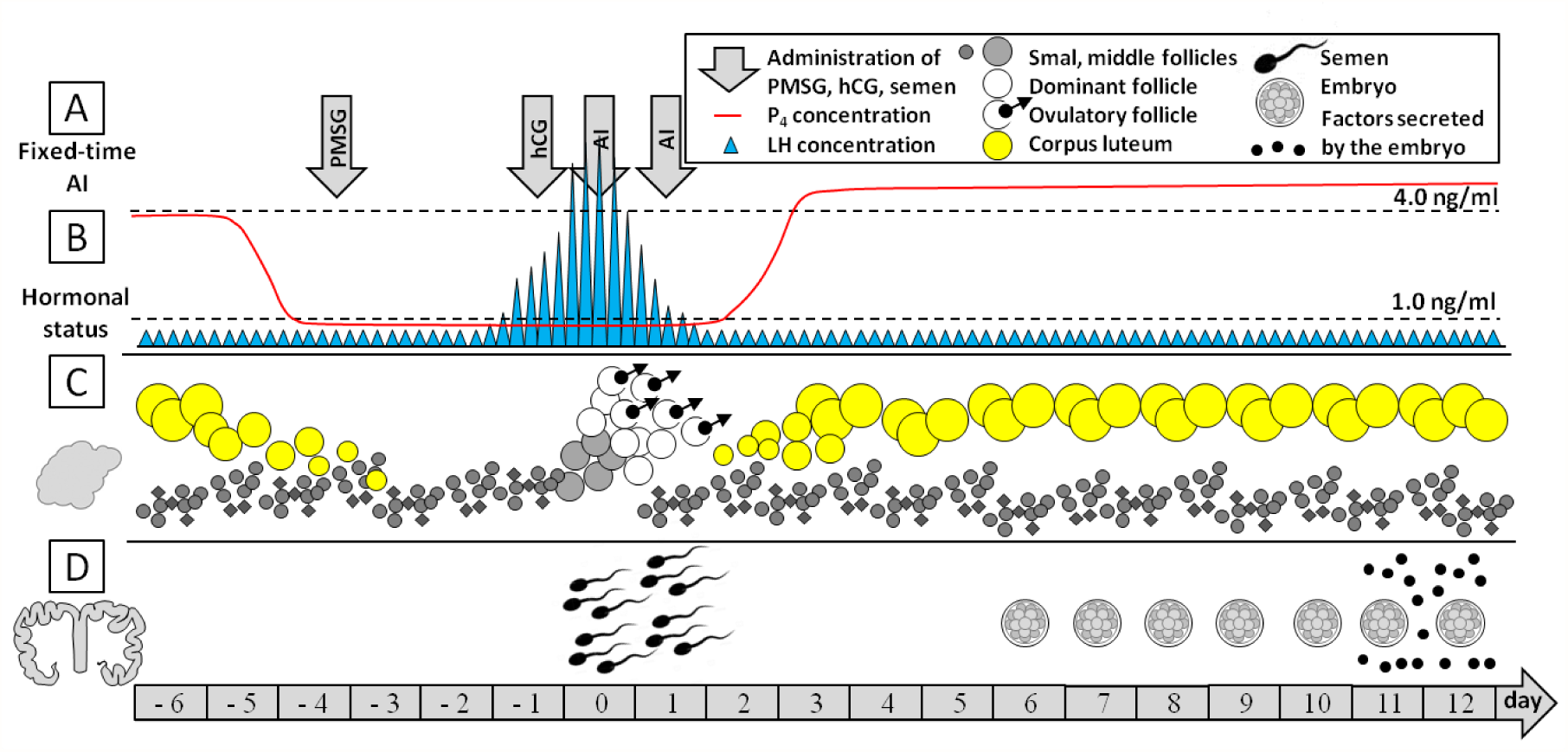
The time-line of considering in: A. procedures of fixed-time artificial insemination, B. hormonal status, C. ovarian status, D. content of uteri lumen.

Day −6 and −5 correspond to the diestrus stage (hormone concentration: LH <1.0 ng/ml; P4 >1.0 ng/ml), whereas days from −4 to 1 show changes in the proestrus (day −4 to −2; LH <1.0 ng/ml; P4 <1.0 ng/ml) and estrus (day −1 to 1; LH >2.0 ng/ml; P4 <1.0 ng/ml) stage of the estrus cycle, respectively. During day −1 the concentration of LH increased (LH >4.0 ng/ml; P4 <1.0 ng/ml) concurrently and the AI were conducted twice (day 0 and 1). Finally, the P4 concentration gradually increased (LH <1.0 ng/ml; P4 >1.0 ng/ml) from day 2 and remained at a high level (LH <1.0 ng/ml; P4 >4.0 ng/ml) since day 12 after AI. At day 12 sows were euthanized (sodium pentobarbital; Morbital 100.0 or more mg/kg b.w., i.v., Fa; Biowet Pulawy, Poland) and the pregnancy has been confirmed after flushing the reproductive tract in order to collect embryos according to Pawlinski et al. (2017). Good quality, viable embryos have been collected from each animal. In both uterine horns 12 - 19 embryos in size 6 - 8 mm were observed under a dissecting microscope (Fa; Olympus SZ51, Japan).

### 2.2. EMG data acquisition

From day 4. after surgery the EMG signal was measured continuously before and during estrus synchronization as well as during AI and early pregnancy. The EMG recording was continued for 12 days after first AI. The EMG signal was acquired with a 3-channel transmitter (TL10M3-D70-EEE; DSI, USA) with sampling frequency - 100 Hz. Obtained analogue signal was digitalized and sent by radio waves to the telemetric receiver (DL10 analogue output; Fa; DSI, USA). The band-pass digital filter (5-50 Hz) and the notch filter were applied for lowering the power line interference from the EMG raw data (PowerLab 4e; Fa; ADInstruments, Australia). Mean and linear trends were removed and the EMG signal was analyzed using LabChart 8 Reader (Fa; ADInstruments, Australia) software. The electrical activity of myometrium reflecting uterine contraction was defined as burst when amplitude exceeded 5 μV for longer than 3 s, the individual bursts were separated with non active periods from longer than 0.5 s, according to Gajewski et al. (2004). In the time domain bursts parameters were determined using duration (D, s), amplitude (A, mV) and root mean square (RMS, mV) of bursts as well as duration of pauses between bursts (P, s). In the frequency domain EMG signal spectral content was calculated after signal multiplication by Hamming window as in Muthuswamy and Thakor (1998) and Domino et al. (2016). By means of the FFT: the dominant frequency (DF, Hz; the frequency of the maximum power within the selected frequency range), maximum power (Max P, V^2^ x10^−11^; the maximum power within the selected frequency range) and minimum power (Min P, V^2^ x10^−24^; the minimum power within the selected frequency range) was assessed for each burst (Fig. 1B). In recent papers the action potentials measured *in vivo* were determined mainly by one statistical outcome variable at a time (A, RMS, D or their univariate statistic) (Gajewski et al., 2004; Garfield and Manner, 2007a; Domino et al., 2016; Pawlinski et al., 2016). Since Garfield and Maner (2007b) and Govindan et al. (2015) suggested analysis of the uterine electrical activity in frequency domain as a potential indicators of predicting uterine contraction, the methods taking into account the effects of all features on the recent responses of interest has become necessary. Therefore the following probabilistic model was introduced including both timer domain and frequency domain features and their mutual interdependency.

### 2.3. EMG data processing

Obtained data are represented the form of matrix X ∈ ℝ^*t*×*n*^ with elements *x*_*j,i*_ where *j* index correspond to realizations and *i* index corresponds to features being univariate marginals. Each day we have selected *t* = 400realizations representing action potential activities (bursts). Such data were obtained manually, using LabChart v.8.0 Reader (ADInstruments, Australia) software, from signals registered in over 120 min time period at all animals. Obviously, order of realization is irrelevant for the statistical analysis. In our approach we took under consideration 7 features (Duration; Pause; Amplitude; RMS; MaxP; MinP; DF), which order is consistent with indexing. The single realisation vector was x_*j*_ =[*x*_*j*,1_,…,*x*_*j*,n_] ∈ ℝ^1×7^.

While introducing the probabilistic model, supposed data are modelled by the multivariate distribution with the following multivariate probability density function PDF *ƒ*(x): ℝ^*n*^ → ℝ. We assumed that probabilistic model does not change during a given day, but may change between days. The characteristic function (Kendall and Stuart, 1946; Lukacs, 1970) of *ƒ*(x) was given by:

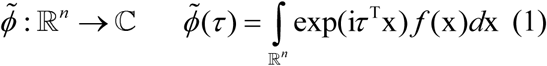

where *τ* is an argument vector *τ* = [*τ*_1_,…,*τ*_*n*_] and i is an imaginary unit. The *d*^*th*^ multivariate cumulant is an object (a tensor) indexed by *i*_1_,…*i*_*d*_, where each *i*_*k*_ ∈ [1,2,…,*n*], with following elements:

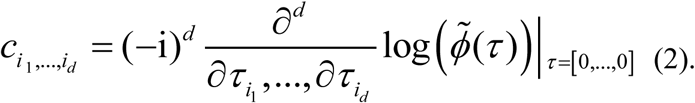

It was easy to show that for the probabilistic model of the multivariate Gaussian distribution 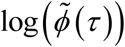 is quadratic in *τ*, hence for *d* ≥ 3 Equation (2) gives zeros. In consistency with a standard notation for *d* ≥ 3 we have higher order cumulants. Observe that for *d* = 1 we have a mean vector and for *d* = 2 a covariance matrix, we call them the standard cumulants.

Following Domino and Gawron (2019) one can use the Frobenius norm on the higher order multivariate cumulant to measure how far data are from hypothetical multivariate Gaussian model. Importantly multivariate Gaussian distribution model does not allow for simultaneous extreme values of many features (Domino et al., 2018b), hence the appearance of such extreme simultaneous events should be reflected in higher order cumulants values. The Frobenius norm of the *d*^*th*^ cumulant tensor can be defined as follows:

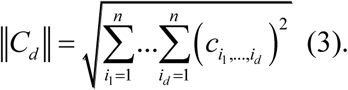

To calculate higher order cumulants we use the efficient algorithm introduced in Domino et al. (2018b) and implemented in `Julia` programming language (Domino et al., 2017b).

#### 2.3.1 Information extraction and hierarchy of features

For each day of the peri-fertilization period we have collected data in the following matrix form: X ∈ ℝ^400×7^. Due to the specifics of the signal collected, values of various features differ by many orders of magnitude. Hence we are rather interested in information carried by the whole features set. To overcome an impact of different orders of magnitudes of features we normalise them by dividing each by its standard deviation. Normalized data 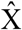 are defined in element wise notation:

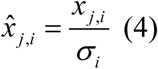

where *σ*_*i*_ is a standard deviation of the *i*^*th*^ column (*i*^*th*^ marginal) of X.

It would be interesting to determine the hierarchy of features including information they are caring. For this purpose we use the Joint Skewness Band Selection (JSBS) method that was primary introduced to analyse information stored in non-Gaussian features of hyperspectral images (Geng et al., 2015). Moreover the JSBS was used for information analysis concerning artificially generated data by means of copulas (Domino and Glos, 2018c). Recall that the application of the JSBS to analyse biomedical data is pioneer. Let us move to the formal introduction of the JSBS.

To introduce formally the JSBS we need to refer to Equation (2) again. We can introduce the 3^*rd*^ cumulant’s tensor *C*_3_ ∈ ℝ^*n*×*n*×*n*^ being an 3 dimensional Array and containing all cumulant’s elements *c*_*i*_1__,_*i*_2__,_*i*_3__, for the formal tensor definition see (Kolda and Bader, 2009). Observe that:

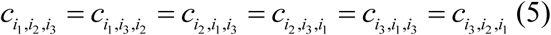

since differentiation is commutative, this observation is called the super-symmetry of the cumulant’s tensor (Schatz et al., 2014). Following (Kolda and Bader, 2009) the first mode unfolding of 3^*rd*^ cumulant’s tensor, noted as (*C*_3_)_(1)_ ∈ ℝ^*n*×*n*^2^^ is the matrix composed of following elements:

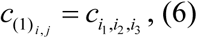

where its second index is:

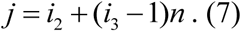

in our case. Reffer to Kolda and Bader (2009) for the general definition. Recall that, there exists other possible methods to unfold a tensor into a matrix. However due to the supper-symmetry of *C*_3_, all those methods are equivalent Kolda and Bader (2009). This is an argument for taking a first mode unfold as its mathematical notation is simple. Having defined a tensor unfold into matrix, we can define the “joint skewness” matrix:

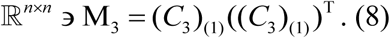

As argued in Geng et al. (2015) such matrix measures information extracted by the 3^*rd*^ cumulant about a non-Gaussian distribution of data. Finally to measure the relative information about the non-Gaussianity, we use the following target function:

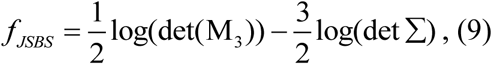

which is a logarithm of the target function introduced in Geng et al. (2015). The log values are better to present, and logarithm is strictly increasing function, hence it does not affect the information hierarchy. Since Σ ∈ ℝ^*n*×*n*^ is a covariance matrix, we may think about *ƒ*_*JSBS*_ as the measure of information carried by the 3^*rd*^ cumulant, but not by a covariance matrix. Referring to Figure 3 the covariance matrix is little affected by the biological process, hence its information is redundant here. Importantly determinant is the product of eigenvalues of the matrix and as such it is proportional to the information hyper-ellipsoid volume (Sheffield, 1985) of the matrix.

**Fig. 3.**
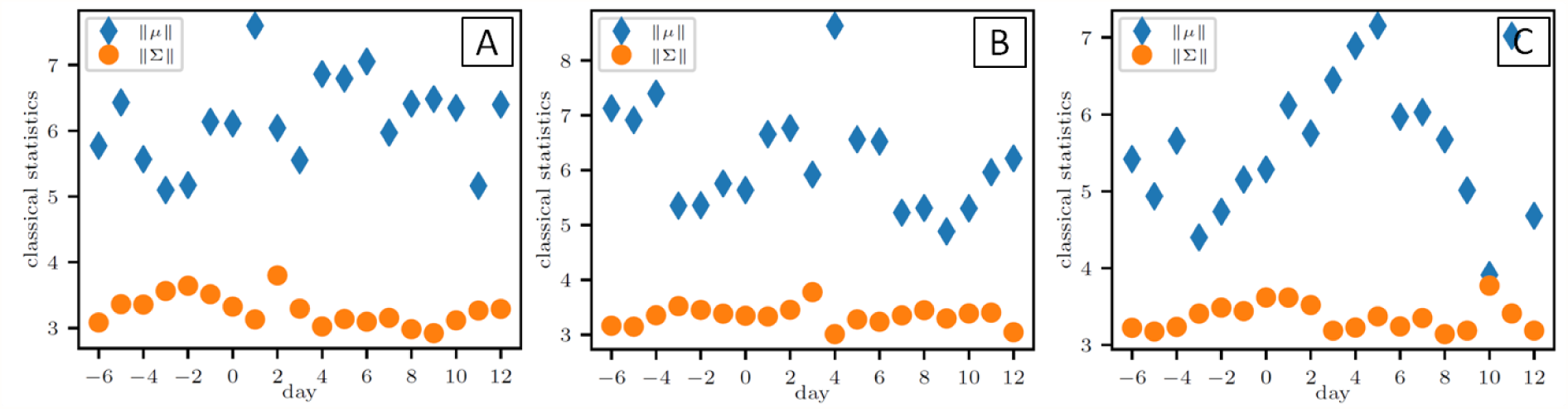
Standard statistics; mean vector (‖*μ*‖) and covariance matrix (‖Σ‖) for seven features of EMG signal obtained in the different parts of uterus from: A. channel 1, B. channel 2, C. channel 3.

Given a target function we perform the following features elimination scheme:

1. we compute the *ƒ*_*JSBS*_ for all data 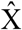,
2. we remove one marginal of 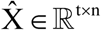 in such a way that the target function of new 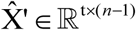 is maximises, removed feature (marginal) is less informative,
3. we repeat *n* – 1 times the procedure in point 2 to obtain the order of features by their information significance.

## 3. Results and discussion

The clear, free of artefacts EMG signal was recorded from all electrodes positioned along one side of uterus from day −6 to day 12 in respect to AI. We used the Frobenius norm on the higher order multivariate cumulant and found that considering together time and frequency features of EMG signal was extremely non-Gaussian distributed as following ‖*C*_3_ ‖ ≫ 0 and ‖*C*_4_‖ ≫ 0 in all points of designed time-line.

Beginning with the standard cumulants the Frobenious norm of the mean vector (for *d* = 1) and the covariance matrix (for *d* = 2) of 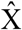 are presented in Figure 3. One can observe that there are no sharp changes as expected for those values in our experimental system. No significant changes are visible and repeatable in all three measurement sites (Ch1, Ch2, Ch3). Such sharp changes would be informative for our system analysis, since we expect the probabilistic model to change between days due to the change of the complex dynamics of EMG signal pattern. Hence such standard multivariate statistics are not informative here.

Following on higher order multivariate statistics the Froebenious norm of the 3^*rd*^ cumulant’s (for *d* = 3) and the 3^*rd*^ cumulant’s (for *d* = 4) of 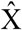, measuring of the distance from the Gaussian multivariate distribution, are presented in Figure 4 and 5, respectively. There black dash lines gives the 0.975 quantile if particular date were from the Gaussian multivariate distribution. The examined features are high above black dash line indicating that the pattern of EMG activity is significantly far from hypothetical multivariate Gaussian model, in all measurement sites repetitively. Our results indicate than bioelectrical signal, such as uterine EMG, brings within much more information than has been recently achieved from univariate statistic (Jedruch et al., 1989a; Gajewski et al., 2004; Pawlinski et al., 2016; Pawlinski et al., 2017) or multivariate Gaussian distribution model (Fig. 3). Secondly there are high jumps (a value of the norm may double) yielding a change of the probabilistic model and suggesting transformations of EMG signal pattern.

**Fig. 4.**
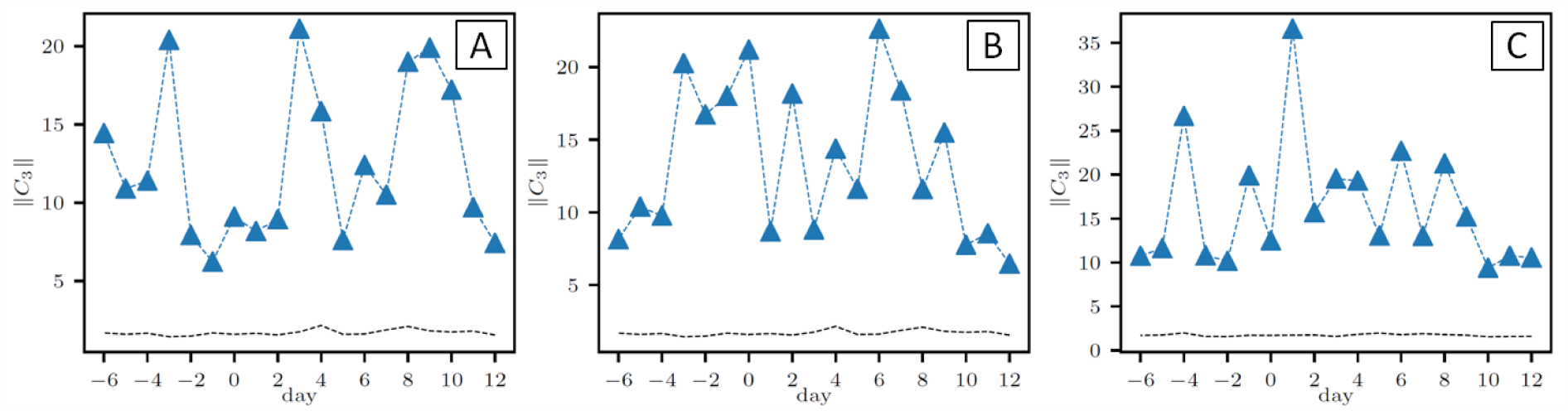
Higher order multivariate statistics; The 3^*rd*^ multivariate cumulant (‖*C*_3_‖) for seven features of EMG signal obtained in the different parts of uterus from: A. channel 1, B. channel 2, C. channel 3. The black dash lines gives the 0.975 quantile if particular date were from the Gaussian multivariate distribution. Footnotes: Black dashed lines are generated numerically, for each day we estimate parameters of the hypothetical multivariate Gaussian distribution from data and perform 250 experiments of sampling *t* realisations from this hypothetical distribution and compution ‖*C*_3_‖ for those hypothetical data.

**Fig. 5.**
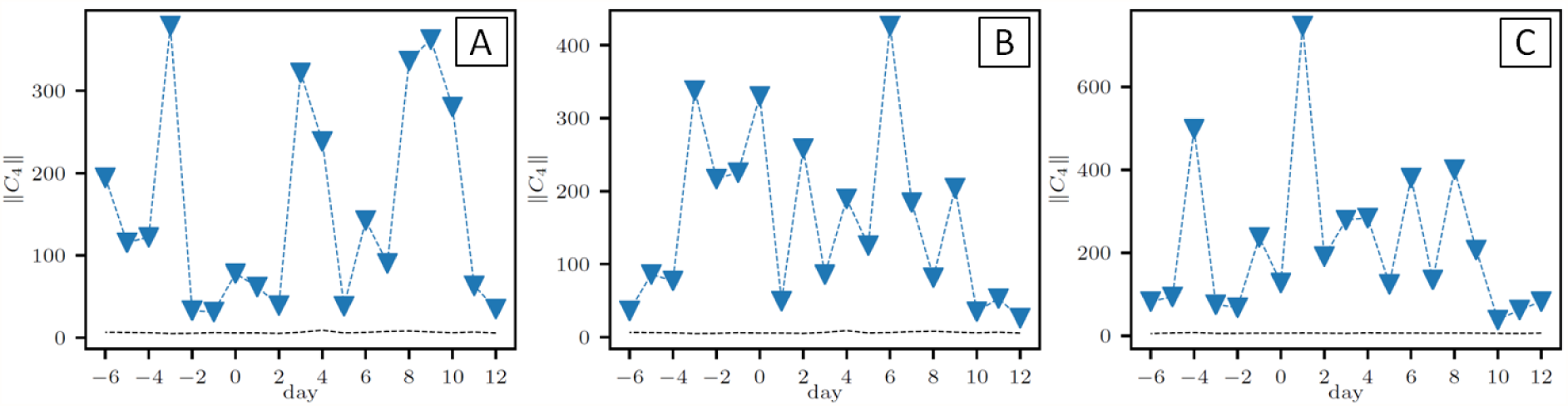
Higher order multivariate statistics; The 4^*th*^ multivariate cumulant (‖*C*_4_‖) for seven features of EMG signal obtained in the different parts of uterus from: A. channel 1, B. channel 2, C. channel 3. The black dash lines gives the 0.975 quantile if particular date were from the Gaussian multivariate distribution. Footnotes: Black dashed lines are generated numerically, for each day we estimate parameters of the hypothetical multivariate Gaussian distribution from data and perform 250 experiments of sampling *t* realisations from this hypothetical distribution and compution ‖*C*_4_‖ for those hypothetical data.

To explain such behaviour we suggest the appearance of simultaneous extreme values of EMG signal features after estrus cycle synchronization beginning in corpus uteri (day −4, Fig. 4C, Fig. 5C) and spreading along uterine horn (days −3 to 0, Fig. 4B, Fig. 5B) into the tip of uterine horn (day −3, Fig. 4A, Fig. 5A). In the middle of uterine horn the most changes in myoelectrical activity pattern between days −3 and 0 have been observed. Garfield and Yallampalli (1994) suggested in uterine muscle signal parameters and its propagation depend on properties of wave propagation medium, which differed significantly witch changes of reproductive status (Jedruch et al., 1989a; Jedruch et al., 1989b; Andersen and Barclay, 1995; Kitazawa et al., 2001; Mueller et al., 2006). Those differences in the properties of the medium apply components of intracellular systems at the level of the SMCs plasma membrane both the number of gap junction and expression of hormone (including reproductive hormones) receptors (Garfield and Maner, 2007a). The changes in hormone (and their receptors) concentration and ion channel distribution interact in dynamical manner to modulate the membrane potential, the type of action potential, and the ionic conductance in the myometrium (Wray et al., 2015). Therefore the changes in probabilistic model may also be the most visible in estrus, when the endogenous regulatory impact is the highest. Our results indicate the largest changes in EMG signal pattern during estrus in the uterine horn, which coincides with the highest receptors for ovarian hormones (Kitazawa et al., 2001; Mueller et al., 2006) and gap junction (Garfield et al., 1977; Andersen and Barclay, 1995) output in this part of reproductive tract.

The second high jump occur shortly after AI again in the first in corpus uteri (day 1, Fig. 4C, Fig. 5C) and then in the middle (day 2, Fig. 4B, Fig. 5B) and in the tip (day 3, Fig. 4B, Fig. 5B) of uterine horn. After AI due to sperm cells motion and myometrium contractions, a small subpopulation of sperm are very quickly (within 1-2 hours) transported from the uterus towards the utero-tubal junction where they colonize sperm reservoir (Rath et al., 2008). The fate of the majority of sperm is elimination and the rapid removal of sperm is thought to prevent acquired immune response against sperm (Hansen, 2011; Katila, 2012). The myometral contractions takes active part in the elimination of sperm, which starts simultaneously with semen transport. Frequent uterine contractions carry semen back and forth in the uterus (Katila et al., 2000; Sumransap et al., 2007). Because the elimination of sperm and post insemination inflammatory products is important for fertility, the efficient myometrial contractions are a prerequisite for uterine cleaning and preparing the intrauterine environment for the development of the embryo (Katila, 2012). The crucial changes in probabilistic model may also be observed.

The third high jump was detected only in uterine horn, both in the tip (days 8 to 10, Fig. 4C, Fig. 5C) and the middle (days 6 to 8, Fig. 4B, Fig. 5B) corresponding to the descent of embryos into the uterine lumen. That intrauterine migration of porcine embryos occurs between day 6. and 10(11). (day 0 = fertilization) (Ziecik et al., 2011). In this period the changes in uterine activity during may be expected. Pope et al. (1982) showed that myometrial contractility increased concomitantly with embryo migration. Similarly Markiewicz et al. (2016) described an increase of contractile activity in the porcine uterus with embryos. The changes of probabilistic model which we are demonstrated are in line with the recent studies.

Interestingly, at days 11 and 12 the evidences of the largest order were observed in corpus and horns of uterus at once (Figs. 4-5). At the day 11 after AI, the embryos starting secretion of estrogen, which stimulates luteoprotective prostaglandin E_2_ (PGE_2_) synthesis in the process of maternal recognition of pregnancy (Ziecik et al., 2011). In this process embryo signals its presence in the uterus and mother receives and accepts this signal (Short, 1969). When embryo signals are systemically recognized by mother, the corpus luteum function and also pregnancy are maintained, therefore the changes in uterine activity pattern during this period should be also expected. The complex system in which billions of cells comprising myometrium, responds similarly and accordingly to biological signals transmitted through uterine luminal epithelium and underlying smooth muscle layers (Dantzer, 1985; Franczak and Bogacki, 2009). The interaction between embryos and uterine wall provides the quiescence of the uterus requiring to maintain pregnancy.

Referring to Figure 4 one can conclude that the 3-order multivariate cumulant gives optimal information about biological phenomena that we are investigate in. The large estimation error of higher order cumulants, especially the 4^*th*^ order, should be associated with the relatively small data samples *t* = 400. This disadvantage of our method may be improved by increasing radically data sample size as one discussed in Appendix A (Domino et al., 2018b). Nevertheless, in our case higher order multivariate cumulants are much more informative that classical statistics. The following cumulants based tools find application to analyse our data. The norm of the higher order cumulants tensors is used to analyse how far the probabilistic model of data is from a Gaussian multivariate distribution. Moreover the 3^*rd*^ cumulant’s tensor based Joint Skewness Band Selection (JSBS) is used to order features according to information they carry (Geng et al., 2015). The hierarchy of features has been successfully determined using the pioneer application of the JSBS to analyse biomedical features. An application of *ƒ*_*JSBS*_ allowed to determinate the dependence structure of EMG patterns with an effective EMG feature. Some of examples of target function values during features elimination process are presented in Figure 6. Sometimes we have a maximum if all features are included, and sometimes after removing one feature as in Table 1. The explanation is not straight forward, since *ƒ*_*JSBS*_ is rather used to compare equally large subsets of features. Recall that the JSBS is searching for a subset of features that has meaningful non-Gaussian joint distribution, while highly non-Gaussian data are informative for our model. We have shown that Min P, Max P and P features carries highest information according to our model, while DF, D and RMS are little informative. Hence we have a combination of features both in time domain and frequency domain that are meaningful.

**Fig. 6.**
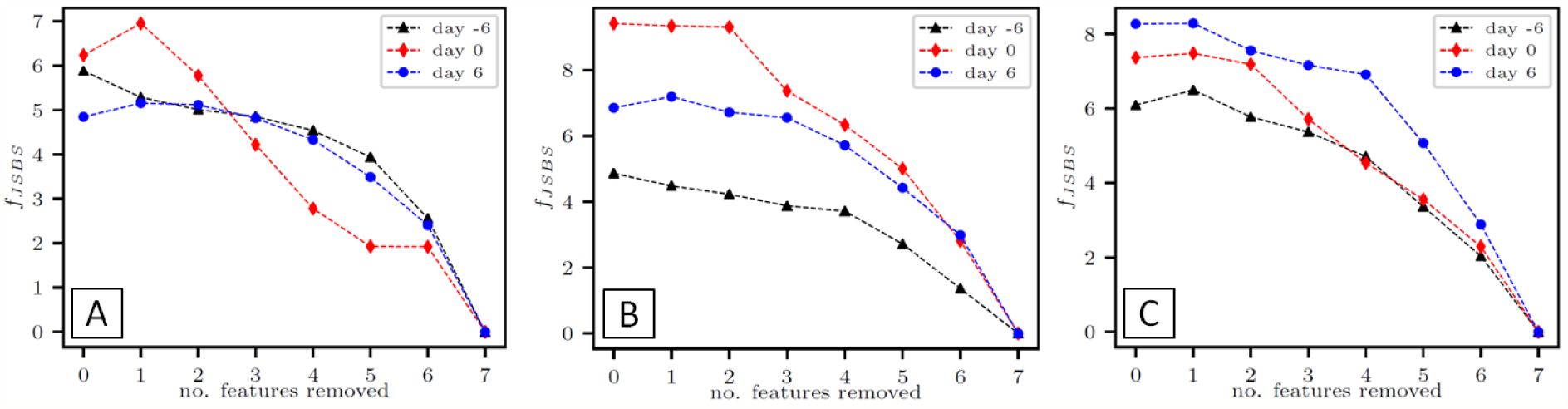
Higher order multivariate statistics; 3^*rd*^ cumulant’s tensor based JSBS (*ƒ*_*JSBS*_); The examples of target function values during features elimination process for seven features of EMG signal obtained in the different parts of uterus from: A. channel 1, B. channel 2, C. channel 3 at day −6, 0, 6 of experiment.

**Table 1.** The hierarchy of EMG signal features in subsequent points of designed time-line of peri-tertilization period.

| **Fetures / day** | **Ch** | **- 6** | **- 5** | **- 4** | **- 3** | **- 2** | **- 1** | **0** | **1** | **2** | **3** | **4** | **5** | **6** | **7** | **8** | **9** | **10** | **11** | **12** |
| --- | --- | --- | --- | --- | --- | --- | --- | --- | --- | --- | --- | --- | --- | --- | --- | --- | --- | --- | --- | --- |
| D | 1 | IV | VI | VI | V | VI | VI | IV | VI | VII | VI | VI | VI | V | IV | VI | VI | VI | IV | III |
| Duration | 2 | IV | IV | IV | VI | VI | VI | VI | VI | VII | V | VI | VI | VI | VI | VII | IV | VI | III | VI |
| [s] | 3 | VI | VI | VI | IV | VI | VII | VI | VII | VII | V | II | VI | IV | VI | VI | VII | VI | VI | VI |
| P | 1 | II | I | II | VI | V | V | VI | II | V | IV | III | I | I | I | II | II | III | II | IV |
| Pause | 2 | I | II | II | IV | I | I | I | I | VI | I | II | I | I | I | I | III | I | II | III |
| [s] | 3 | III | III | I | VI | V | I | I | I | III | II | VI | I | II | I | II | II | V | I | II |
| A | 1 | VI | V | IV | III | IV | IV | II | V | III | II | II | V | VI | VI | V | V | V | VI | VI |
| Amplitude | 2 | V | V | VI | III | V | IV | V | V | III | VI | IV | IV | V | IV | V | VI | IV | V | IV |
| [mV] | 3 | IV | IV | IV | III | III | V | V | V | IV | IV | IV | IV | V | III | IV | IV | II | IV | V |
| RMS | 1 | VII | VI | V | IV | III | III | III | VI | II | V | IV | IV | IV | V | IV | IV | IV | V | V |
| Root mean square | 2 | VI | VI | V | V | IV | III | VI | VI | VI | III | V | V | IV | V | IV | V | V | VI | V |
| [mV] | 3 | V | V | V | V | II | IV | IV | IV | VII | VI | III | III | VI | IV | V | V | I | V | IV |
| Max P | 1 | III | III | III | II | I | II | V | III | I | III | V | III | III | III | III | III | II | III | II |
| Maximum power | 2 | III | III | III | II | III | V | III | II | II | VI | III | III | III | III | III | II | III | IV | II |
| [V<sup>2</sup>] | 3 | II | II | III | II | IV | III | III | III | II | III | V | II | III | II | III | III | III | III | I |
| Min P | 1 | I | II | I | I | II | I | I | I | IV | I | I | II | II | II | I | I | I | I | I |
| Minimum power | 2 | II | I | I | I | II | II | II | III | I | II | I | II | II | II | II | I | II | I | I |
| [V<sup>2</sup>] | 3 | I | I | II | I | I | II | II | II | I | I | I | V | I | V | I | I | IV | II | III |
| DF | 1 | V | VII | VII | VII | VII | VII | VII | VII | VI | VII | VII | VII | VII | VII | VII | VII | VII | VII | VII |
| Dominant frequency | 2 | VII | VII | VII | VII | VII | VII | VII | VII | V | VII | VII | VII | VII | VII | VII | VII | VII | VII | VII |
| [Hz] | 3 | VII | VII | VII | VII | VII | VI | VII | VI | V | VII | VII | VII | VII | VII | VII | VI | VII | VII | VII |

Footnotes: D, P, A, RMS, Max P, Min P, DF - the EMG signal features considered in experiment; Ch - the different parts of uterus from (channel 1, channel 2, channel 3); day - the days considered in experiment; I - VII - the hierarchy of features indicated by 3<sup>rd</sup> cumulant's tensor based JSBS.

Our results obtained in animal *in vivo* model supporting the hypothesis of a functional role of uterine activity in promotion of fertilization both in estrus (days −4 to 0) and after AI (days 1), preparation of uterine environment to embrace the embryos (days 2 to 3) and support of embryos migration during early pregnancy (days 6 to 10). We suspect the middle of uterine horn as the most heavily involved area affecting the phenomena of embryo development and implantation and the Min P, Max P and P features as the most informative in the model of spontaneous myoelectrical activity of complex SMCs systems in peri-fertilization period. Due to the fact that we obtained good quality, viable embryos at day 12, the studied development of early pregnancy should be considered proper. The applied approach is suitable in recognition of crucial stages in peri-fertilization processes providing successful fertilization and early pregnancy maintenance.

## 5. Conclusions

An additional novelty of this paper resides in the porcine EMG signal analyzing methodology based on the use of higher order multivariate cumulants. We have applied those cumulants to study the dependence structure of EMG patterns with an effective EMG feature, and then have built up the myoelectrical activity templates for recognizing complex uterine contraction according to crucial stages for successful fertilization and early pregnancy maintenance.

Based on our results, we may conclude that due to non-Gaussian joint distribution of features such as, higher order multivariate statistics such as cumulants, have to be used to determine the pattern of myoelectrical activity in reproductive tract. Using those tools, we confirmed the expectance that the probabilistic model changes on a daily base, while analyzing the complex biological system of porcine SMCs and the functional role of uterine activity in peri-fertilization period. We speculate the middle of uterine horn and the Min P, Max P and P features are crucial elements of the uterine activity model, whereas further information is needed to elucidate the coordination and propagation of EMG signal. Concluding, the higher order multivariate cumulants modelling offers a unique opportunity to understand the mechanisms underlying uterine contractility and development of early pregnancy in animals. The authors belief that similar dynamics may appear in human beings due to high applicability of porcine preclinical models.

## Acknowledgements

This work was conducted in the Veterinary Research Centre WULS (WCB) and the Center for Biomedical Research (CBB) supported by EFRR RPO WM 2007-2013. Additionally, KD acknowledge the support of the National Science Centre, Poland under project number 2014/15/B/ST6/05204.

## Conflict of interest

None.

